# Programming aliphatic polyester degradation by engineered bacterial spores

**DOI:** 10.1101/2024.07.16.603759

**Authors:** Ziyu Cui, Masamu Kawada, Yue Hui, Seunghyun Sim

## Abstract

Enzymatic degradation of plastics is a sustainable approach to addressing the growing issue of plastic accumulation. The primary challenges for using enzymes as catalysts are issues with their stability and recyclability, further exacerbated by their costly production and delicate structures. Here, we demonstrate an approach that leverages engineered spores that display target enzymes in high density on their surface to catalyze aliphatic polyester degradation and create self-degradable materials. Engineered spores display recombinant enzymes on their surface, eliminating the need for costly purification processes. The intrinsic physical and biological characteristics of spores enable easy separation from the reaction mixture, repeated reuse, and renewal. Engineered spores displaying lipases completely degrade aliphatic polyesters and retain activity through four cycles, with full activity recovered through germination and sporulation. Directly incorporating spores into polyesters results in robust materials that are completely degradable. Our study offers a straightforward and sustainable biocatalytic approach to plastic degradation.

## INTRODUCTION

Plastic products are indispensable to our modern society and enable many comfortable aspects of our daily lives, ranging from food packaging to bullet-proof vests. Compared to hard materials such as metals and ceramics, they possess numerous practical characteristics – low weight, economical mass production, flexibility, and modularity in mechanical and biological properties. Because their explosive demand and production accompanied our economic growth in the last decades, plastic waste can be found everywhere and continues to accumulate in our environment.^1, 2^ According to the recent modeling study performed by Rochman and colleagues, increased waste management capacity alone cannot compensate for the projected growth in plastic waste generation, and major technological innovation is imperative for handling this impending crisis.^3^

Some synthetic plastics (e.g., poly(lactic acid) (PLA)) contain hydrolyzable moiety, such as ester, and can be broken down upon spontaneous hydrolysis. However, spontaneous hydrolysis of these plastics typically requires a high temperature, harsher conditions, and a long time to be completed.^4^ For example, biodegradable plastics poly(ε-caprolactone) (PCL) and PLA are indifferentiable in landfills, and even thermophilic digesters operating at 48–60 °C cannot yield full breakdown of these plastics.^5^ Physical recycling is the currently predominant method.^6-8^ In this process, collected used plastics are shredded into flakes, sorted, cleaned, and melted into a new batch of recycled materials. The mechanical stress and photo-oxidation caused during the physical recycling process generally decrease the durability of the recycled product.^6^

Enzyme catalysis offers a unique solution to these challenges as a green method to facilitate the plastic degradation process.^9^ Several plastic materials have been shown to be degraded by enzymes. For example, a number of fungal/bacterial lipases, esterases, hydrolases, and cutinases are shown to degrade aliphatic or aromatic polyesters.^10, 11^ Embedding functional enzymes within polyester to program degradation of materials has been explored by a number of studies.^12-16^ Typically a protecting agent, such as random heteropolymers or polymeric coatings, is used to preserve the enzyme function during harsh processing conditions involving high temperatures or organic solvents.^12-14^ Despite the apparent merits of enzymes in chemical manufacturing processes in terms of mild reaction environment, specificity, fast reaction rate, and biocompatibility,^6, 17, 18^ using enzymes for a wide range of applications in a scalable and economical manner remains challenging. In addition to the costly production process to obtain them in high purity, enzymes are inherently fragile and not reusable. Subjecting them to conditions favorable for typical chemical reactions, such as common organic solvents or high temperatures,^18^ leads to denaturation of their folding structures and subsequent loss of function.^19, 20^

Immobilization of enzymes on solid supports has been shown to effectively alleviate these limitations and is considered essential for the commercial viability of installing enzyme biocatalysts in large-scale production.^21^ The heterogeneous nature of immobilized enzymes allows recycling by simple filtration or centrifugation. Moreover, increased operational stability in high temperatures and other harsh conditions is often observed in immobilized enzyme systems.^21, 22^ Typically, enzymes are immobilized onto solid support via chemical bond formation or physical entrapment. Immobilization of enzymes on bacterial spores features additional benefits by leveraging the autonomous biological process of protein synthesis and assembly on the spore surface and affording a straightforward harvesting process for spores via centrifugation or filtration. In particular, spores can tolerate environmental stresses common in industrial settings, such as high temperatures, nutrient deprivation, and organic solvents.^23^ Spores can survive these abiotic conditions and germinate into vegetative cells when nutrients become available.^23, 24^ To harness the unique physical and biological characteristics of spores as a heterogenous scaffold for enzyme immobilization, we have developed the T7 RNA polymerase-enabled high-density protein display (TIED) platform to display 10^6^–10^7^ recombinant proteins on bacterial spores.^25^ These TIED spores displaying recombinant enzymes exhibited robust catalytic activities, resilient in harsh conditions, and enabled recycling and complete renewal of catalytic activities through the cycle of germination and sporulation.^25^

In this project, we explore the utility of engineered bacterial spores to program degradation of aliphatic polyesters with two complementary strategies (Figure 1). The spores are engineered to display *B. subtilis* lipases through the TIED platform. First, we showed that exogenously added engineered spores can completely degrade a range of aliphatic polyester films: PCL, PLA, and poly(lactic-co-glycolic acid) (PLGA). These aliphatic polyester films were completely and uniformly degraded within 2 days by simply incubating them with the engineered spores in aqueous solutions at moderate temperatures. Secondly, we demonstrate the direct incorporation of these catalytic spores, without any protecting agent, into aliphatic polyester to yield degradable composite materials with comparable physical properties to pristine polyester materials. They undergo spontaneous degradation by a simple incubation in aqueous solutions. Furthermore, we showed that spores retained catalytic activity when recycled multiple times, and upon attrition of activity, it can be fully renewed upon germination and subsequent sporulation.

**Figure 1.**
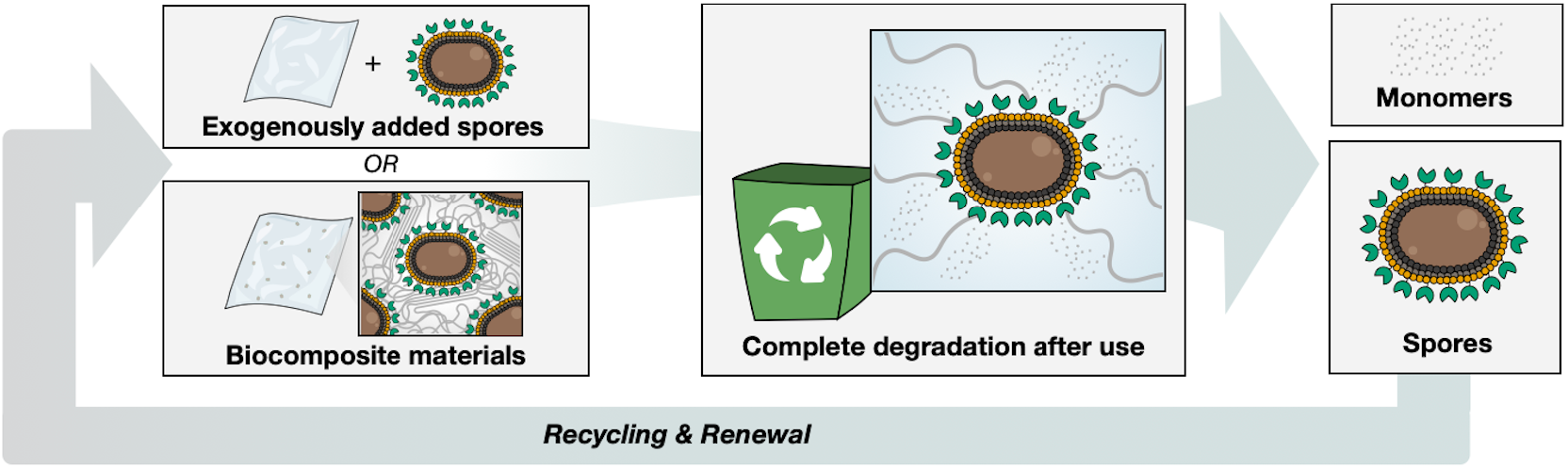
Schematic illustration of two approaches presented in this work for degrading the polyesters by engineered bacterial spores.

## RESULTS

### Complete degradation of PCL films by engineered spores

The engineered spores displaying *B. subtilis* Lipase A and *B. subtilis* Lipase B through the TIED exhibited high enzyme loading density and robust small molecule hydrolysis activity.^25^ Based on their activities, we selected the promoter and fusion partner combinations of *P*_*cotV*_ and CotZ for Lipase A (TIED-LipA) and *P*_*cotG*_ and CotZ for Lipase B (TIED-LipB) for our study. PCL films (6.25 cm^2^, 5.0 mg ± 0.5 mg) incubated in the spore-containing solution (1 mL OD_600_ 0.5 solution supplemented with 800 *μ*L 100 mM Tris-HCl, pH 8.0) were degraded completely over time (Figure 2a). Scanning Electron Microscopy (SEM) images revealed that compared to the intact films (Figure 2b), degrading PCL films have rough surface textures with holes and cracks (Figure 2c). The weight of the PCL films decreased upon incubation with TIED-LipA (Figure 2c, Supplementary Figure S1A) and TIED-LipB (Figure 2d, Supplementary Figure S1B). films, but not with wild-type spores, which have endogenous lipase and other hydrolases on the surface (Figure 2e, Supplementary Figure S1C), even with 100-h incubation. The same concentration of free-floating Lipase A and Lipase B (0.5 mg mL^-1^), based on the western blot-based quantitation of enzymes on TIED-LipA and TIED-LipB,^25^ completely degraded the PCL films in 4 h (Supplementary Figure S2A and S2B). This result shows that free-floating enzymes show faster hydrolysis kinetics for heterogeneous substrates such as PCL film than spore-immobilized ones. Notably, PCL crystallinity was maintained at around 55% in the course of enzymatic degradation by TIED-LipA, TIED-LipB, and their free-floating counterparts (Figures 2c and 2d, Supplementary Figure S1). Taken together, our data shows that engineered spores high-density of hydrolytic lipase displayed on their surface show robust, complete, and uniform degradation of PCL films, including crystalline region.

**Figure 2.**
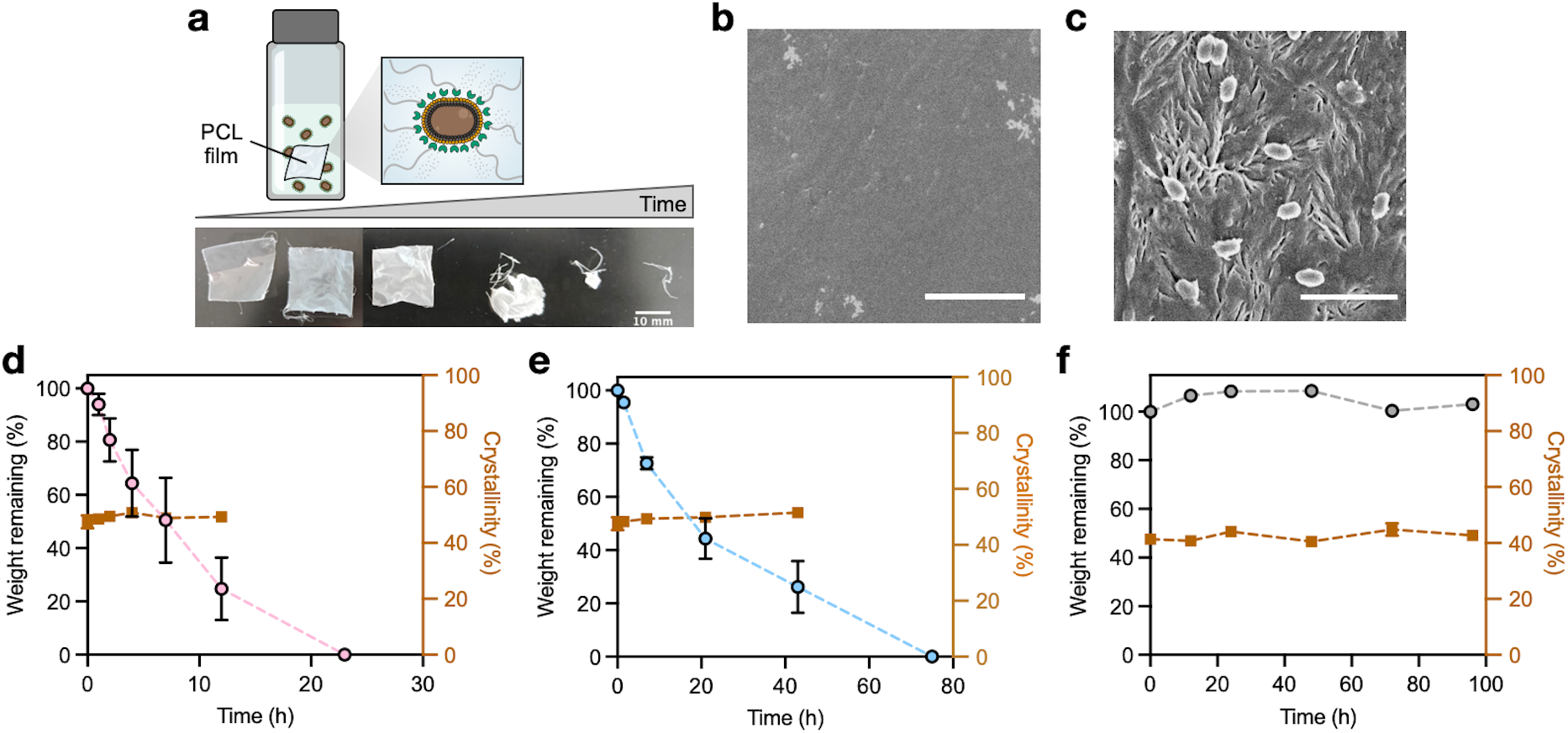
PCL film degradation by engineered spores. (a) Schematic representation of the experimental setup and photographic images of PCL films incubated with spore suspension over time. (b, c) A representative scanning electron microscope (SEM) image of (b) a pristine PCL film and (c) a PCL film collected during the degradation process. Scale bars 5 *μ*m. (d–f) Mass (circles) and crystallinity (brown squares) of the remaining PCL upon incubation with (d) TIED-LipA, (e) TIED-LipB, and (c) wild-type PY79 spore suspension. The reaction conditions for all experiments are identical (42 °C, 800 rpm). *N* = 3 biological replicates, and error bars are standard error of means (s.e.m.).

### Complete depolymerization of the semi-crystalline aliphatic polyester into monomers

Motivated by the complete degradation of added PCL films, we set out to investigate the molecular composition and distribution of the degrading PCL fragments. We collected all PCL fragments present in the system by freeze-drying the entire reaction mixture at different time points and extracting PCL fragments with tetrahydrofuran (THF) (Figure 3a). The molecular weight and distribution of PCL fragments were analyzed by gel permeation chromatography (GPC). The average molecular weight of PCL was generally unchanged during the degradation (Figures 3c and 3d, Supplementary Tables 1–3).

**Figure 3.**
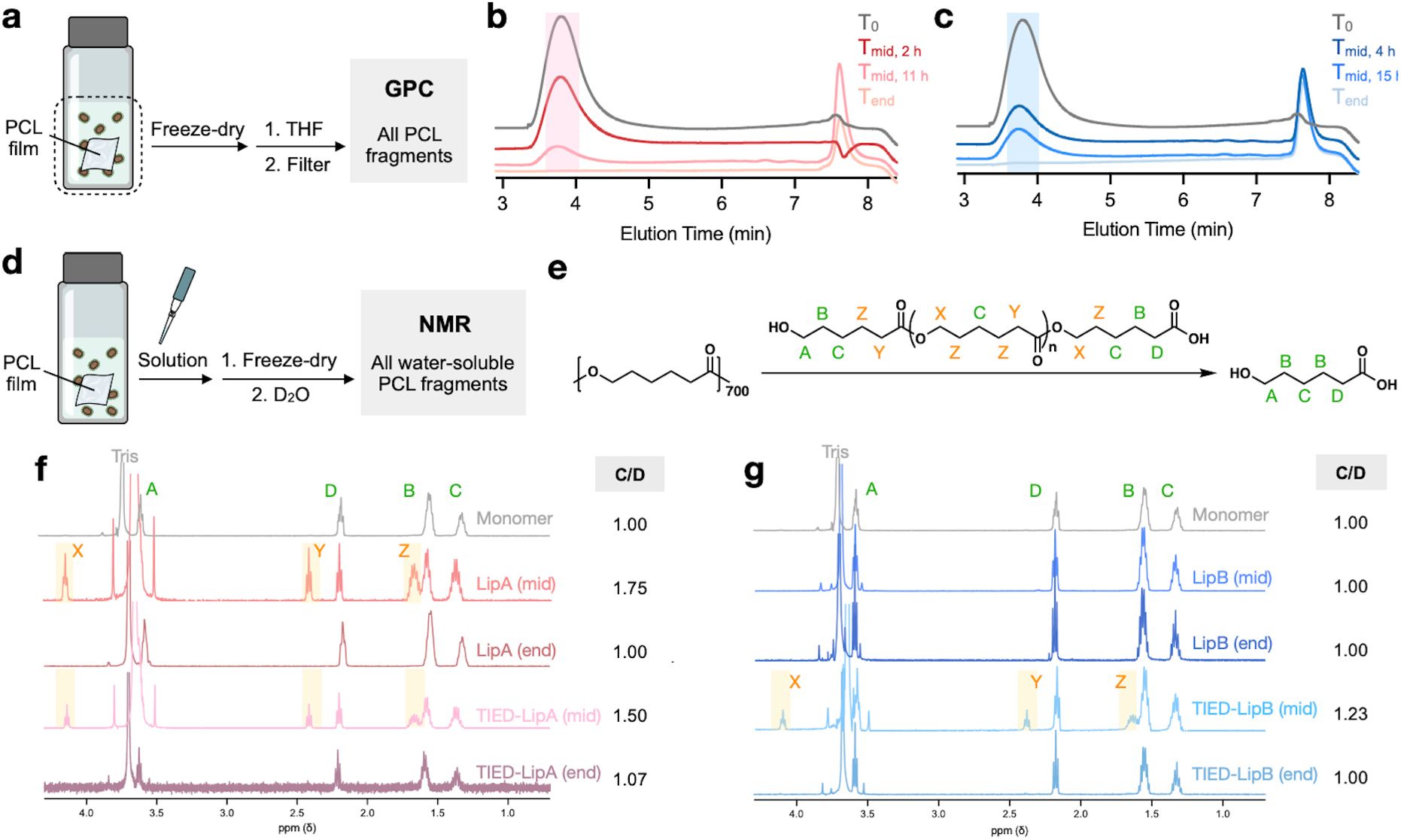
TIED-LipA completely depolymerizes PCL into monomers. (a) Schematic illustration of the gel permeation chromatography (GPC) sample preparation. (b, c) GPC trace of all PCL fragments before and after incubation with (b) TIED-LipA and (c) TIED-LipB. (d) Schematic illustration of the nuclear magnetic resonance (NMR) sample preparation. (e) General reaction scheme of PCL hydrolysis. (f, g) ^1^H NMR spectra (D_2_O) of water-soluble PCL fragments incubated with free-floating (LipA and LipB) and spore-displayed enzymes (TIED-LipA and TIED-LipB). The gray curve corresponds to monomer (6-hydroxyhexanoic acid) spectra taken in the same condition (D_2_O, 293.1K).

We then set out to analyze water-soluble fragments by ^1^H NMR during the degradation. The aqueous supernatant from the reaction mixtures was collected, lyophilized, and dissolved in Deuterium Oxide (D_2_O) (Figure 3d). Upon PCL degradation, the number of chain ends increases, and their adjacent protons become distinguishable by chemical shifts (Figure 3e, Supplementary Figure 5). Peak C represents a common set of methylene protons at the gamma position, while Peak D is unique to the chain end. Therefore C/D value should represent the average number of repeating units. The monomer spectrum shows a C/D value of 1.00 (Supplementary Figure 4). The endpoint was defined as when there were no visible PCL fragments, and the midpoint for each sample was set to be the time point where 50% weight loss of PCL films was expected (Figures 2c and 2d). In all of the midpoints of the reaction mixtures, C/D values were between 1 and 2. Peaks X, Y, and Z show the presence of oligomers as peak X represents the methylene protons adjacent to the ester, which are not present in the monomer form, and peaks Y and Z are distinguishable from Peaks D and B due to differences in chemical environments in oligomers and monomers. The peak assignments were further validated by the spectrum of pristine PCL in CDCl_3_ (Supplementary Figure 3). In all of the endpoints of the reaction mixtures, C/Ds are 1.00 or very close to 1.00 (Figures 3f and 3g, Supplementary Figures 6–13), demonstrating that the PCL was completely hydrolyzed into monomeric form in the degradation process. Considering the relative bulkiness of enzymes, especially the spore-immobilized ones, we speculate a classical surface erosion process liberating short-chain PCL fragments to be the main mode of action for the complete degradation of PCL films into monomers. Dissolved short-chains would then be rapidly hydrolyzed into monomers by enzymes in solution. Our data also shows that immobilizing enzymes on the spore surface does not alter the main mode of degradation for PCL.

### Expanding the aliphatic polyester substrate scope of TIED-LipA

In addition to PCL, poly(D,L-lactic acid) (PDLLA) and poly(lactic-co-glycolic acid) (PLGA) were chosen to test the generality of TIED enzymes for hydrolyzing aliphatic polyesters because of their wide use in the biomedical engineering and 3D printing.^26, 27^ PDLLA and PLGA films were majority amorphous, and their enthalpy peaks did not show significant change while the weight loss was observed (Supplementary Figure S14). Both PDLLA and PLGA completely degraded upon incubation with TIED-LipA spores (Figures 4a and 4d), and it generally took about 30 hours for completion (Figures 4b and 4e, Supplementary Figure S15). PDLLA and PLGA films incubated with wild-type spores also exhibited about ∼20% mass reduction over 100 hours (Figures 4b and 4e), showing that these amorphous polyesters are more susceptible to hydrolysis by a small number of hydrolytic enzymes naturally present on the spore surface. Scanning Electron Microscopy images revealed that degrading PDLLA (Figure 4c) and PLGA (Figure 4f) films have rough surface textures with holes and cracks compared to the pristine films (Figures 4c and 4f). Similar to PCL (Figure 2), these results demonstrate that engineered spores effectively degrade a wide range of aliphatic polyesters.

**Figure 4.**
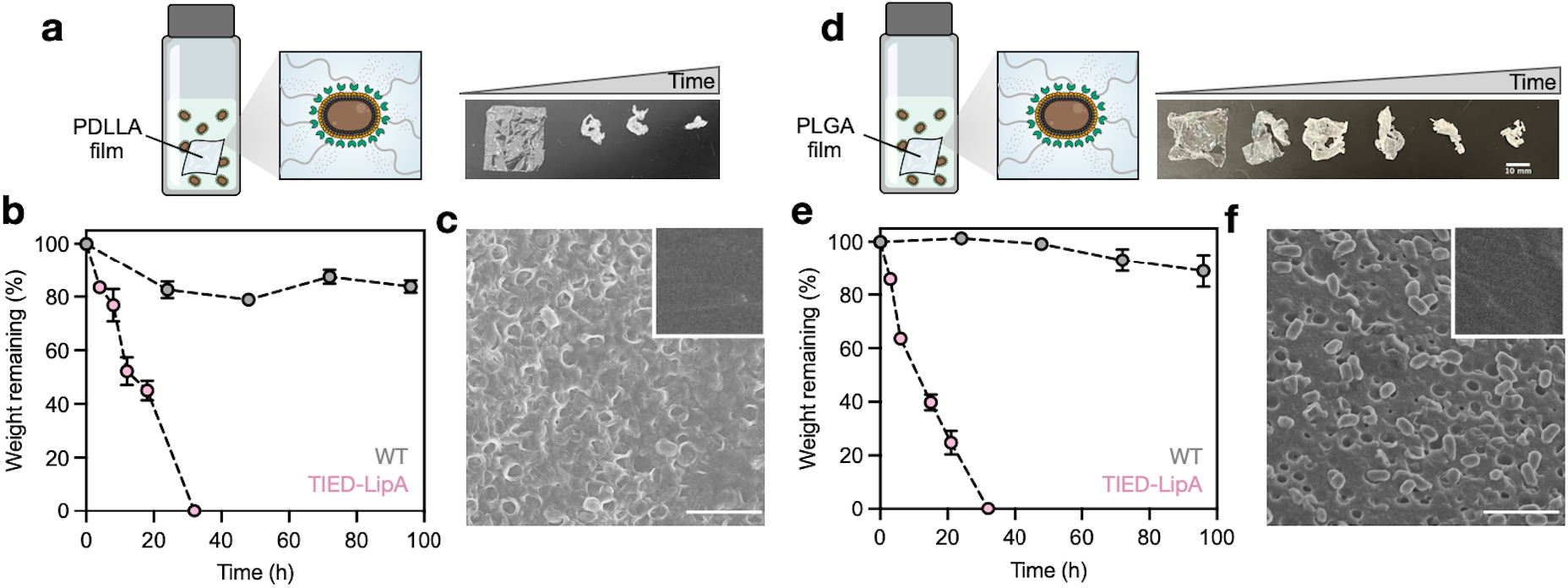
Degradation of Poly(D,L-lactic acid) (PDLLA) and poly(lactic-co-glycolic acid) (PLGA) films. (a) Schematic representation of the experiment setup (left) and photographic images of PDLLA films incubated with TIED-LipA spore suspension over time (right). (b) Mass of the remaining PDLLA films upon incubation with TIED-LipA (pink) or wild-type spore (gray) suspension. (c) A representative SEM image of a PDLLA film collected during the degradation process. The inset is a representative SEM image of a pristine PLGA film. Scale bar 5 *μ*m. (d) Schematic representation of the experiment setup (left) and photographic images of PLGA films incubated with TIED-LipA spore suspension over time (right). (e) Mass of the remaining PLGA films upon incubation with TIED-LipA (pink) or wild-type spore (gray) suspension. (f) A representative SEM image of a PLGA film collected during the degradation process. The inset is a representative SEM image of a pristine PLGA film. Scale bar 5 *μ*m. *N* = 3 biological replicates, and error bars are standard error of means (s.e.m.).

### Recycling and renewal of catalytic spores for PCL degradation

One of the unique advantages of the spore display system is that (1) catalyst particles can be easily separated from the reaction mixture through filtration or centrifugation, and (2) upon attrition of catalytic activities, a fresh batch of spores can be regenerated by germination and re-sporulation. We set out to test recyclability and renewability in the context of plastic degradation with PCL and TIED-LipA. Suspension of TIED-LipA was treated with fresh PCL films every 24 hours. TIED-LipA after each reaction, after collecting any remaining PCL films, were washed and subjected to a new reaction (Figure 5a). Collected pieces of PCL film were weighed to calculate the percentage of mass loss during each cycle, which reflects enzymatic activity within 24 hours. After the first cycle, we observed a gradual attrition of activity in the following cycles (Figure 5b). This result shows that while the recycled spores retain catalytic activities, they tend to be slower than the pristine ones. After the 4th cycle, the spores were collected from the reaction mixture and germinated into vegetative cells, which yielded a new batch of catalytic spores under a standard sporulating condition.^25^ Gratifyingly, the renewed spores completely recovered their original activity and achieved complete degradation of the PCL film within 24 hours.

**Figure 5.**
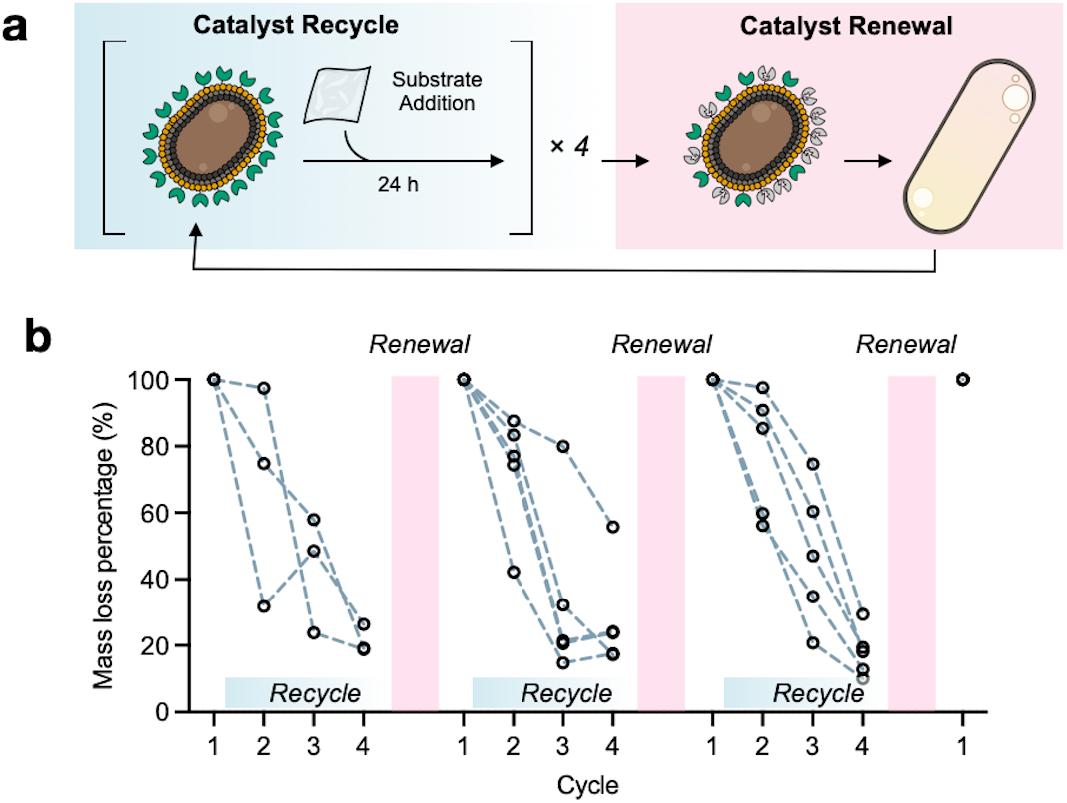
Recycling and renewal of catalytic spores. (a) Schematic illustration of recycling and renewal of catalytic spores. (b) Recycling and renewal of TIED-LipA spores. Each reaction cycle lasts 24 hours. *N* = 3 (biological replicates) in the first recycle and *N* = 5 (biological replicates) in the following renewal and recycling experiments.

### Programming spontaneous degradation of biocomposite materials by directly incorporating the engineered spores

Previous studies have examined strategies to embed enzymes into materials to program their spontaneous degradation after use. In most cases, due to the fragile nature of enzymes, a protective agent, such as random homopolymers or polymer coatings, was required to incorporate them into polymeric materials,^12-14^ as the typical processing conditions involve organic solvents or high temperatures. Despite these recent developments, the recoverability of enzymes remains a critical challenge. Given the enhanced resilience of enzymes immobilized on the engineered spores,^25^ we set out to test whether the direct incorporation of catalytic spores into polymeric materials enables spontaneous degradation. Briefly, 12.5 g PCL dissolved in dichloromethane (3 w/v%) was mixed with lyophilized TIED-LipA spore in a powder form (from 1 mL of spore solution in OD_600_= 0.5) and then cast to yield biocomposite materials (Figure 6a). The solvent and concentration of spores were chosen based on the uniformity of spores in the resulting materials (Supplementary Figure S16). SEM images revealed that while the surface of the biocomposite materials was smooth (Figure 6b), intact spores were embedded inside the materials (Figure 6c). The mechanical properties of PCL without engineered spores (Figure 6d) and the biocomposite materials (Figure 6e) were probed by tensile tests. Key physical properties, such as Young’s modulus and toughness, of these two were generally comparable (Figures 6f and 6g), whereas the materials’ ability to elongate (Figure 6h) was slightly reduced upon addition of spores.

**Figure 6.**
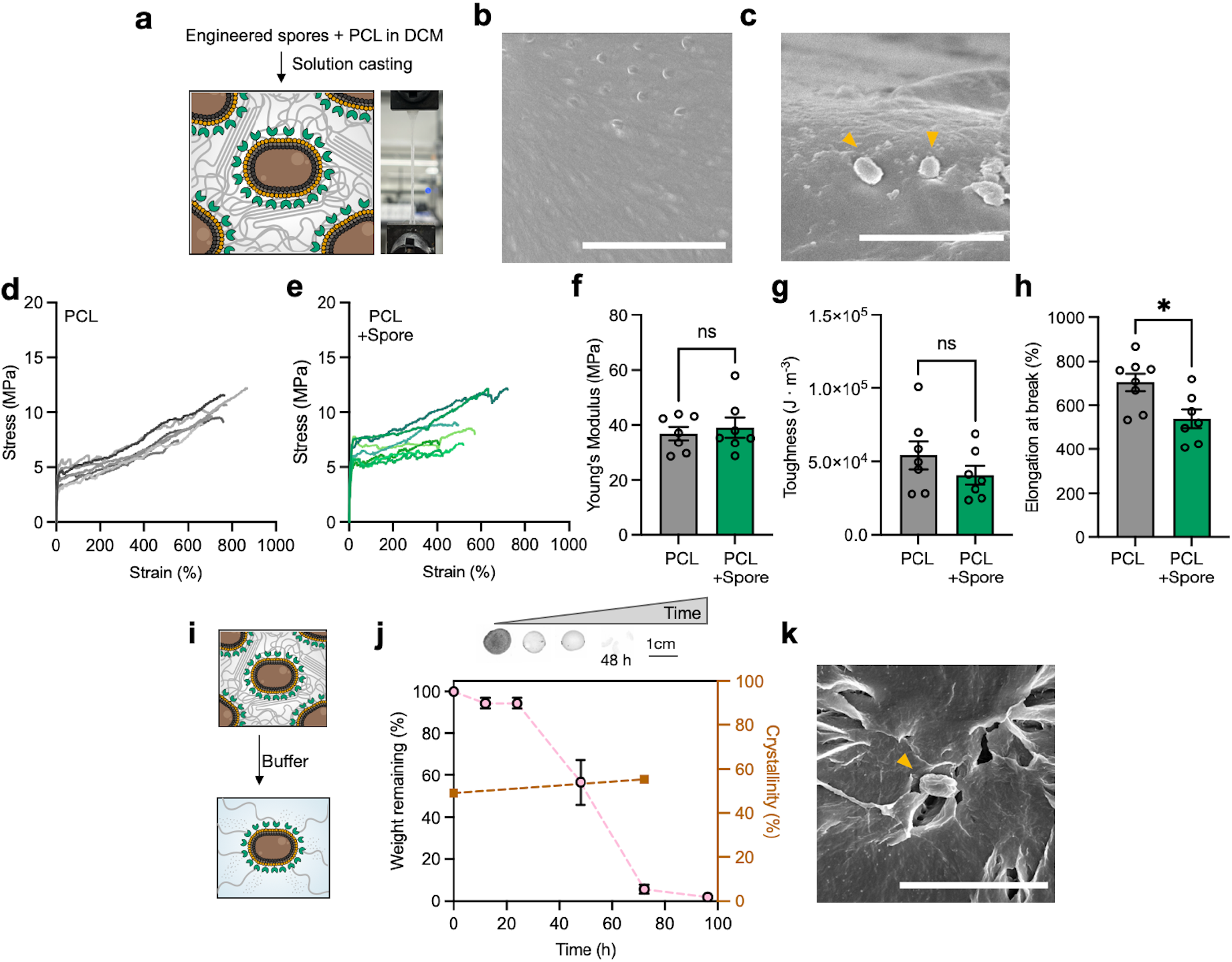
Spontaneously degradable biocomposite materials incorporating catalytic spores. (a) Schematic illustration of biocomposite materials incorporating engineered spores. PCL dissolved in dichloromethane (DCM) was mixed with TIED-LipA and cast to yield biocomposite materials in this study. (b) A representative SEM image visualizing the surface of biocomposite materials. (c) A representative SEM image visualizing the cross-section of biocomposite materials. Yellow arrows indicate embedded TIED-LipA spores. (d, e) Stress-strain curve of PCL (d) with and (e) without TIED-LipA. (f–h) Comparison of (f) Young’s modulus (g) toughness, (h) elongation at break of biocomposite materials and PCL without spores (control). *N* = 7 for manufacturing replicates. (i) Schematic illustration of the spontaneous degradation behavior of biocomposite materials in an aqueous buffer. (j) Images change of biocomposite materials (top). Mass (circles) and crystallinity (brown squares) of the self-degrading biocomposite materials upon incubation (bottom). The images on the top show the progression: pristine materials, 12 hours, 24 hours, and 48 hours of immersion, from left to right. (k) A representative SEM image of degrading biocomposite materials upon immersion into Tris-HCl buffer. The yellow arrow indicates TIED-LipA spores on the degrading material surface. Scale bar 5 *μ*m. *P* values were determined by unpaired two-tail *t*-tests. \**P* < 0.05, \*\**P* < 10^−2^, \*\*\**P* < 10^−3^, \*\*\*\**P* < 10^−4^.

The spontaneous degradation behavior of the biocomposite materials was first tested by immersing them in a buffer solution (1.8 mL Tris 100 mM, pH 8.0) (Figure 6i). The materials degraded into small pieces that were visible but too small to be retrieved within 48 hours (Figure 6j). We acknowledge unavoidable measurement errors at time points before 48 hours due to this reason. By 96 hours, the material becomes completely invisible and disintegrates into solution. SEM images of the degrading materials showed roughened texture and spores on their surface (Figure 6k). The number of viable spores in the buffer solution increased significantly between 24 and 48 hours (Supplementary Figure S17B), during which a significant degradation of the materials was observed (Figure 6j, Supplementary Figure S17A). Collected spores from the materials retained their hydrolytic activity, as demonstrated by their ability to degrade fresh PCL films (Supplementary Figure S17C), albeit with lower activities than the pristine spores. The crystallinity of the biocomposite material was similar to the pristine PCL films and slightly increased over time during the degradation process (Figure 6j). Immersing these materials in an LB medium also resulted in complete degradation (Supplementary Figure S18A) as well as the germination of spores into vegetative cells. The wild-type spores, despite the explosive increase in the number of cells and their general hydrolytic activities, did not result in degradation. Notably, PCL films immersed with TIED-LipA in cellular form maintained their weight. These results illustrate the importance of engineered spores through TIED in achieving complete degradation of aliphatic polyester.

## DISCUSSION

Despite the apparent merits of enzymatic degradation of plastics, especially in terms of sustainability, their fragility and the costly purification processes have been the limiting factors. Spores engineered to display high-density recombinant enzymes present a new solution to this challenge because of the scalable, straightforward, and economical process of producing and isolating catalytic particles. Our work proposes two different strategies with engineered spores with high-density of hydrolyzing enzymes displayed on their surface: (1) degrading aliphatic polyester by exogenously adding engineered spores and (2) incorporating engineered spores directly into materials to program self-degradation properties. We observed the complete degradation of a wide range of aliphatic polyesters by engineered spores. Furthermore, the heterogeneous nature of spore particles enables recyclability, and their robustness enables direct incorporation of catalysts into polymeric materials without stabilizing reagents. Importantly, these catalytic spores can be fully renewed to regain their original activities by germination, cellular growth, and resporulation, which is not possible for chemical catalysts or chemically immobilized enzymes.

Many studies have shown a circular economy of materials by recycling polymeric components.^28-31^ To the best of our knowledge, our work presents the first example of recycling and renewing catalyst components for degrading polymeric materials. Although immobilization can limit the effective collision between the substrate and enzyme, resulting in slower kinetics, engineered spores are capable of degrading semicrystalline polyester completely and uniformly to yield monomeric species in the majority. The self-degradable biocomposite materials presented in our study open the possibility of fine-tuning the lifetime of plastic materials with engineered spores. The physical resilience of the engineered spores will simplify the production process of materials of this kind.

Our results unlock new ways to utilize enzymes to address the plastic issue. The biological programmability, capability to germinate, physical robustness, and heterogeneous nature of *B. subtilis* spores directly address current issues around enzymatic degradation efficiency, scalability, and economic feasibility. We also anticipate that a wide variety of hydrolytic enzymes can be displayed on the spore surface, providing feasible strategies for degrading other types of plastics, such as aromatic polyester or polyolefin. Together, we envision this and the following works in engineered spores will establish sustainable solutions to combat plastic pollution.

## METHODS

### Spore preparation

A single colony on the LB agar plates of wild-type and mutant *B. subtilis* strains (TIED-LipA and TIED-LipB) was inoculated in LB medium (50 mL fresh LB medium in a 250 mL Erlenmeyer flask). Cells were cultured with agitation at 37 °C and 250 rpm until they reached an optical density at 600 nm (OD_600_) of 0.5. Sporulation was then induced using the resuspension method.^32^ Cells were harvested by centrifugation (4000 g, 10 min), and the cell pellets were resuspended in an equal volume of SM medium (0.046 mg FeCl_2_, 4.8 g MgSO_4_, 12.6 mg MnCl_2_, 535 mg NH_4_Cl, 106 mg Na_2_SO_4_, 68 mg KH_2_PO_4_, 96.5 mg NH_4_NO_3_, 219 mg CaCl_2_, 2 g L-glutamic acid, and 20 mg L-tryptophan, pH 7.1). The resuspended cells were transferred back to Erlenmeyer flasks and cultured for 14 h with agitation at 37 °C and 250 rpm. The sporulating cells were collected (4000 g, 10 min) and treated with 50 μg mL^-1^ lysozyme (Sigma-Aldrich, L6876) in phosphate-buffered saline (PBS, pH 7.2, Gentrox, 30-025). The harvested spores were washed with PBS three times, and their morphology was confirmed using an ECHO Revolve Microscope with a phase-contrast objective before use. **Plastic film preparation**. PCL (80 kDa, 440744), PDLLA (75–120 kDa, P1691), and PLGA (50–75 kDa, 430471) pellets were purchased from Sigma Aldrich and used without further purification. The PCL pellets were dissolved in toluene at 20 wt% concentration. The PDLLA and PLGA pellets were dissolved in chloroform at 10 wt%. The polymer films were produced by spin-coating the polymer solutions on a 5-inch wafer. The films were further dried under a vacuum overnight to remove any residual solvent and cut into squares with side lengths of 2.5 cm. Each piece of film was weighed before degradation tests.

### Degradation study with exogenously added spores

Each film was immersed in spore solutions (1.8 mL, OD_600_ 0.278) in Tris-HCl (100mM, pH 8.0) with agitation (42 °C, 800 rpm). The films presented in the degradation process were collected as much as possible at each time point. The collected pieces were immersed and gently washed in 50 mL distilled water for 10 seconds, and the excess water was removed by tapping them with a laboratory-grade wiper (Kimwipes) before drying the films under ambient temperature and pressure. The completely dried films were subjected to weight measurements or differential scanning calorimetry (DSC, DSC 2500, TA Instruments) for measuring crystallinity. For DSC, around 2-4 mg of the PCL films were pressed into aluminum pans and heated from 0 °C to 120 °C at a scan rate of 10 °C min^−1^, PDLLA and PLGA films were scanned from 0 °C to 200 °C. The PCL crystallinity percentage was calculated by normalization of the sample’s enthalpy of melting with the enthalpy of melting for 100% crystalline PCL (151.7 J g^−1^).^33^ The GPC (APC system, Waters) samples were prepared by freeze-drying the entire reaction mixture, extracting organic components with 1 mL THF, and filtering out the non-dissolvable components such as salts, spores, or enzymes using 0.2 *μ*m PTFE filter (Fisherbrand, 09-719G). The injection volume was 10 *μ*L, and ACQUITY APC XT columns (125, 200, 450 2.5 μm) with RI detector were used to analyze molecular weights and dispersity of remaining PCL fragments. The NMR samples were prepared by collecting the aqueous supernatant after centrifuging the reaction mixture at 21,000g for 10 minutes. The collected supernatants were lyophilized and re-dissolved in Deuterium Oxide (D_2_O). As a positive control, monomer (6-hydroxyhexanoic acid, Thermo scientific 1191-25-9) dissolved in Tris buffer (100mM) was collected analogously. The pristine PCL polymer dissolved in CDCl_3_ served as a negative control.

### Recycling and renewal of TIED-A spore

TIED-LipA spores were suspended (OD_600_ = 1) in 100 mM Tris-HCl buffer (pH 8.0). 1 mL of spore suspension with a 6.25 mm^2^ PCL film, 800 uL buffer in 2.0 mL microcentrifuge tubes. The reaction proceeded for 24 hours at 42 °C with agitation (800 rpm) for each cycle. After 24 hours, any remaining pieces of the PCL films were collected, washed with 50 mL distilled water for 10 seconds, and dried under a vacuum overnight. The dried film was weighed to calculate the mass loss percentage. The spores from the reaction mixture were then collected by centrifugation (5000 g, 10 min). The same volume of the Tris-HCl buffer was subsequently added to resuspend the spores. A new piece of the 6.25 mm^2^ PCL film was added and incubated for the next 24 hours. For renewal of TIED-LipA spores, 100 *μ*L of spore suspension after the last reaction was plated on an LB agar plate supplemented with chloramphenicol (5 *μ*g mL^-1^) and spectinomycin (100 *μ*g mL^-1^). Single colonies from the place were inoculated into 5 mL LB to start each culture. Spores were prepared according to the method described above. Degradation reaction conditions were kept constant.

### Preparation and characterization of biocomposite materials

Spore suspensions (OD_600_ = 0.5) were prepared in distilled water and subsequently lyophilized. 12.5 mg PCL (in 3 w/v% DCM) was mixed with lyophilized spore powder from 1 mL of the spore suspension. 10 s vortexing was applied to ensure a homogeneous mixing of spore particles. 100 *μ*L of the mixed solutions was directly drop-cast on the glass substrate to form biocomposite materials. For mechanical testing, 200 *μ*L of the mixed solutions was directly drop-cast into a dog-bone-shape mold (WLH, PTFE). The residual solvents were evaporated under ambient temperature and pressure overnight. For tensile tests, the dog-bone-shaped materials were loaded on the Instron (3365 Universal Testing System) and pulled with a speed of 10 mm min^-1^. For spontaneous degradation tests, circle-shaped biocomposite materials were immersed into 100 mM Tris-HCl buffer with agitation (42 °C, 800 rpm). The remaining materials were collected as much as possible at each time point, washed with 50 mL distilled water for 10 seconds, imaged under Chemidoc MP imaging system, and dried under an ambient environment before measuring crystallinity and SEM imaging.

### Scanning Electron Microscopy imaging

SEM images were acquired from the FEI Magellan 400 XHR SEM at the UC Irvine Materials Research Institute (IMRI). All materials are sputter-coated with 5nm Iridium. Images are acquired under immersion mode, dwell time 3.0 μs, integrated by 12 scannings.

### Statistical analysis

All experimental data were plotted as mean ± s. e. m. unless otherwise mentioned. Statistical significance and *P* values were derived from two-tailed t-tests.

## Supporting information

ESI

## ACKNOWLEDGEMENTS

This work is supported by start-up funds from UC Irvine, and the National Institute of General Medical Sciences of the National Institutes of Health under Award Number R35GM150770. The content is solely the responsibility of the authors and does not necessarily represent the official views of the National Institutes of Health. This research was partially supported by the National Science Foundation Materials Research Science and Engineering Center program through the UC Irvine Center for Complex and Active Materials (DMR-2011967). The authors acknowledge the use of facilities and instrumentation at the UC Irvine Materials Research Institute (IMRI) supported in part by the National Science Foundation Materials Research Science and Engineering Center program through the UC Irvine Center for Complex and Active Materials (DMR-2011967). Nuclear Magnetic Resonance measurements were performed in the NMR facility in the Department of Chemistry, UC Irvine. The authors thank Drs. Xiaofeng Liu, Hyuna Jo, and Ze-Fan Yao for discussions. The authors also acknowledge the help from Yi Liu with SEM and Leslie Liu with GPC. The authors thank the Guan lab at UC Irvine for the use of GPC and the Ardoña lab at UC Irvine for the use of spin coater.

## AUTHOR CONTRIBUTIONS

Z.C. and S. S. designed experiments. Z.C. and M. K. performed and analyzed the NMR data. Z.C. performed all the other experiments and analyzed the data. S.S. and Z.C. wrote the manuscript. S.S., Y.H., and Z.C. conceptualized this work.

## COMPETING FINANCIAL INTERESTS STATEMENT

The authors declare no competing financial interests.

## Notes

### Competing Interest Statement

The authors have declared no competing interest.

