## Supplementary material for "Programming aliphatic polyester degradation by engineered bacterial spores": ESI

### Table of Contents

1. Supplementary Methods ........................................................................................ S2
2. Supplementary Figures 1–18.................................................................................... S3
3. Supplementary Tables 1–3 ..................................................................................... S18
4. References ............................................................................................................. S19

### 1. Supplementary Methods

#### 1.1. Colony forming unit (CFU) from spores released from the material.

Solutions have been vortexed and aliquot to 100 $\mu$L after the film was fished out for characterization. Serial dilution was performed on the aliquot. Dilution was performed to $10^4$-fold dilution from the beginning time point and $10^6$-fold dilution at the latest time point. 10 $\mu$L of each dilution was directly dropped on an LB plate with chloramphenicol (5 $\mu$g mL$^{-1}$) and spectinomycin (100 $\mu$g mL$^{-1}$) supplements to ensure only TIED spores would grow on the plate. Incubated the plates at 37 °C for 16 hours and counted the colonies. The colonies must be individually seen. One-fold concentrate than the ideal dilution, the ideal dilution, and one-fold dilute than the ideal dilution was repeated 3 times. The number of cell-forming units is average from the valid replicates.

#### 1.2. Biocomposite materials degradation in LB.

The circle-shaped biocomposite materials (prepared with TIED-LipA and wild-type spores) were prepared (see Methods). Each film was immersed into a 1.5 mL LB medium in a cell culture tube and incubated at 37 °C, 250 rpm. The LB medium was supplemented with chloramphenicol (5 $\mu$g mL$^{-1}$) and spectinomycin (100 $\mu$g mL$^{-1}$) in the case of biocomposite materials prepared with TIED-LipA to prevent contamination. The remaining materials were collected as much as possible at each time point, washed with 50 mL distilled water for 10 seconds, and dried under an ambient environment before measuring weight.

#### 1.3. PCL degradation by TIED-LipA cell in LB.

Three films were immersed in a 5 mL LB medium supplemented with chloramphenicol (5 $\mu$g mL$^{-1}$) and spectinomycin (100 $\mu$g mL$^{-1}$) to ensure only TIED strains could grow (37 °C, 250 rpm). One single colony of TIED-LipA was introduced to the mixture. Every 12-16 hours, the solutions were refreshed to a fresh LB medium to provide nutrients for cell growth and minimize spontaneous sporulation process. No morphology change was observed from visual inspections. The remaining films were collected and then washed in 50 mL distilled water for 10 seconds, and the excess water was removed by tapping them with a laboratory-grade wiper (Kimwipes) before drying the films under ambient temperature and pressure. The weight at each time point was collected.

### 2. Supplementary Figures

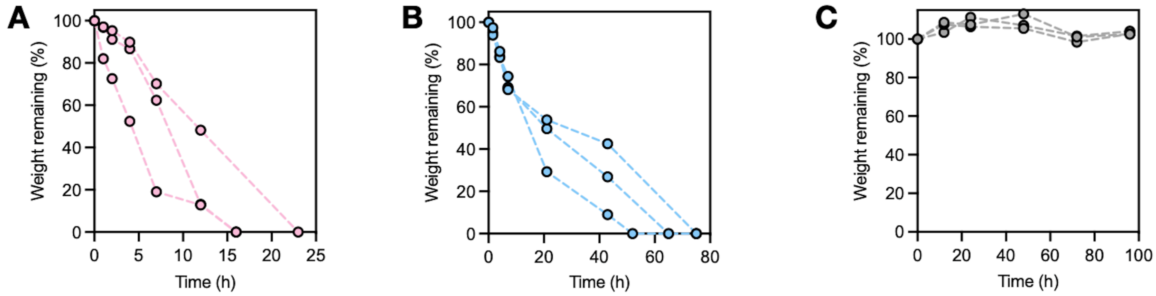


**Supplementary Figure 1. PCL film degradation by spores.** (A–C) Mass percentage of the remaining PCL upon incubation with (A) TIED-LipA, (B) TIED-LipB, (C) wild-type spore suspension (1 mL $OD_{600}$ = 0.5, 800 ). All the mixtures were heated at 42 °C and 800 rpm. Each circle represents individual mass data.

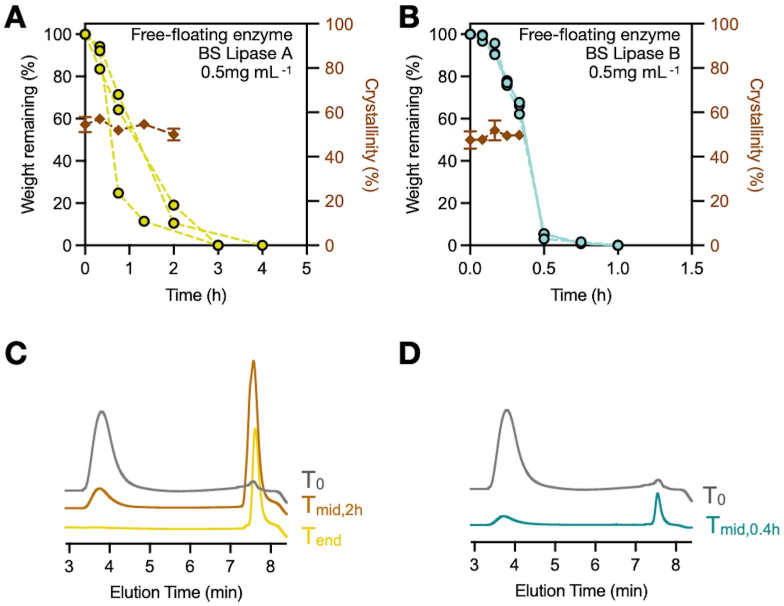


**Supplementary Figure 2. PCL film degradation by free-floating enzyme.** (A,B) Mass (circles) and crystallinity (brown squares) percentage of the remaining PCL upon incubation with 1.8 mL of free-floating (A) Lipase A and (B) Lipase B (0.278 mg $mL^{-1}$), All the mixtures were heated at 42 °C (800 rpm). Each circle represents one individual mass data. Crystallinity data, $N = 3$ experimental replicates, and error bars are standard errors of means (s.e.m.). (C) GPC of all the PCL fragments before and after incubation with free-floating Lipase A. (D) GPC of PCL film before and after incubation with free-floating Lipase B. Midpoint injection volume is 20% of the injection volume of $T_0$.

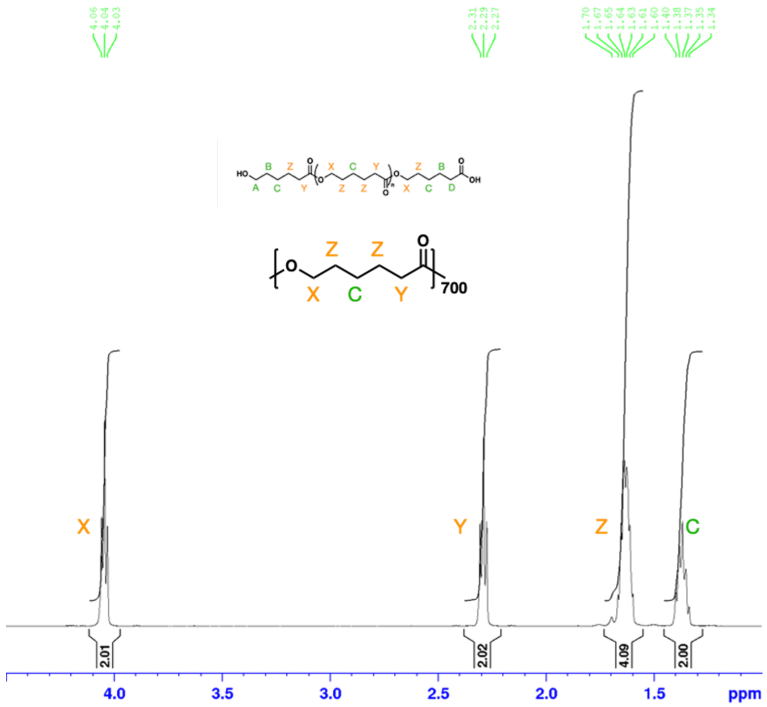

**Supplementary Figure 3. $^1$H NMR ($CDCl_3$) Spectra of PCL.** As peak D was buried with other peaks in the spectra, the C/D value could not be calculated. From the molecular weight, the number of repeating units should be around 700.

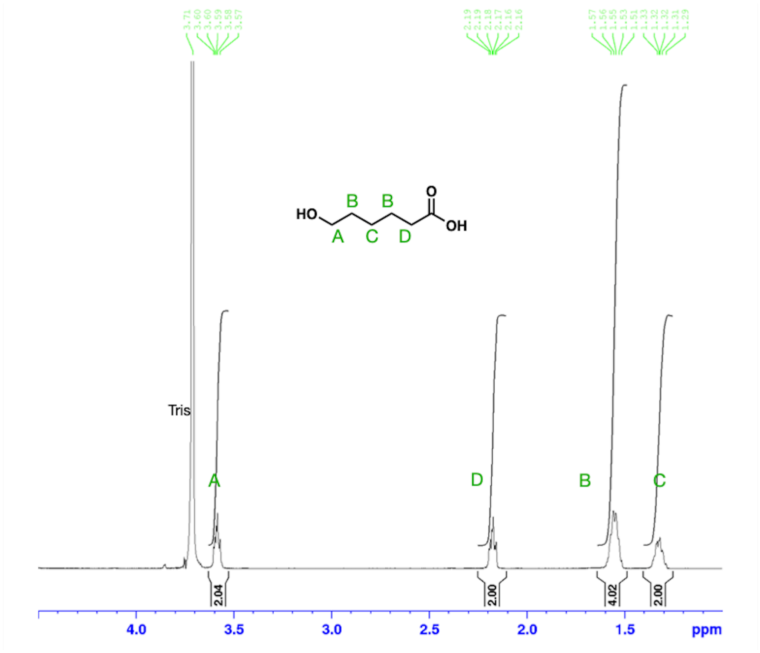

**Supplementary Figure 4. $^1$H NMR ($D_2O$) Spectra of 6-hydroxyhexanoic acid.** The powder was dissolved in 100mM Tris-HCl buffer to mimic the degradation environment. The solutions were lyophilized and subsequently dissolved in $D_2O$ for $^1$H NMR analysis. Spectra was obtained using Bruker instrument GN500 at 500 MHz, referenced to 4.79 ppm Deuterium Oxide peak.

A

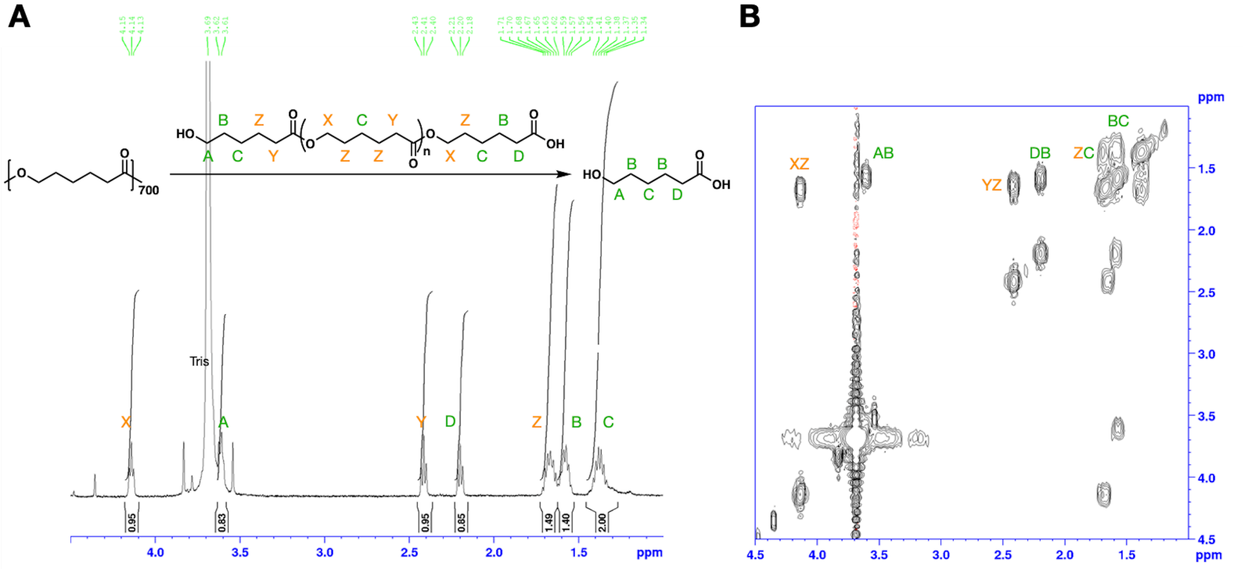


C

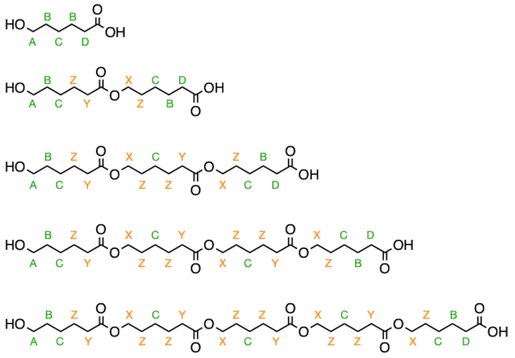

| # | A | B | C | D | X | Y | Z | C/D |
|---|---|---|---|---|---|---|---|---|
| 1 | 2 | 4 | 2 | 2 | 0 | 0 | 0 | 1 |
| 2 | 2 | 4 | 4 | 2 | 2 | 2 | 4 | 2 |
| 3 | 2 | 4 | 6 | 2 | 4 | 4 | 8 | 3 |
| 4 | 2 | 4 | 8 | 2 | 6 | 6 | 12 | 4 |
| 5 | 2 | 4 | 10 | 2 | 8 | 8 | 16 | 5 |
| ... | ... | ... | ... | ... | ... | ... | ... | |
| n | 2 | 4 | 2n | 2 | 2n-2 | 2n-2 | 4n-4 | n |

**Supplementary Figure 5. $^{1}$H NMR analysis of mixtures containing oligomers and monomers.** (A) $^{1}$H NMR spectrum of a PCL degradation mixture containing oligomers and monomers in $D_2O$. (B) $^{1}$H-$^{1}$H COSY NMR spectrum of the midpoint mixture. (C) Peak assignments for potential PCL hydrolysis products: units starting from 1 (monomer) to 5 (pentamer). The table shows calculations for total peak intensity.

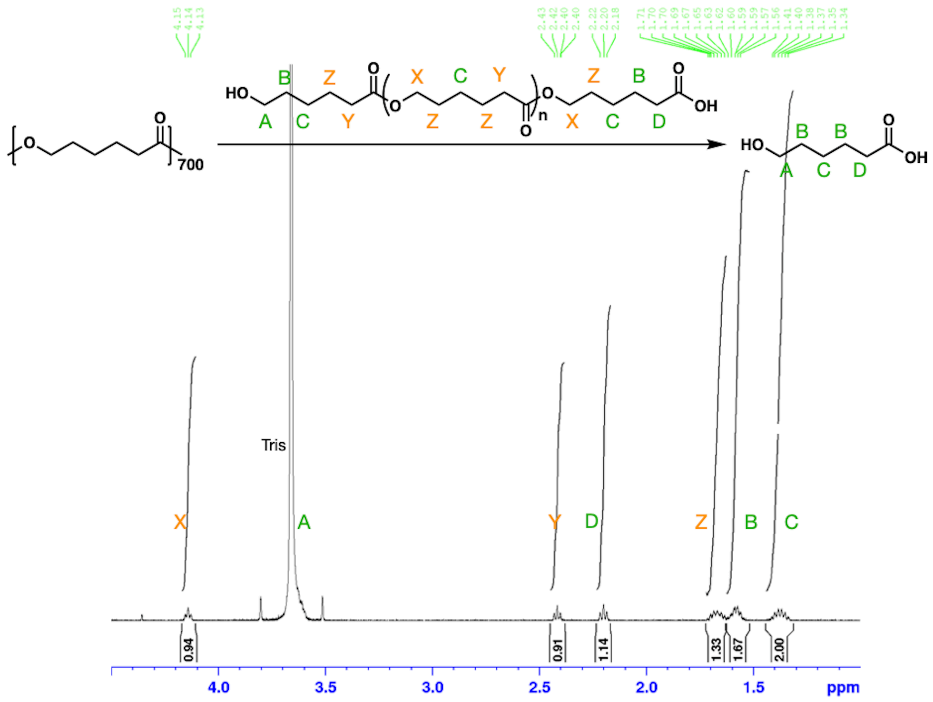


| C/D |
|---|
| 1.75 |

**Supplementary Figure 6.** **$^{1}$H NMR *(*$D_2O$*)* Spectra of the midpoint mixture collected from the PCL degradation process with free-floating enzyme Lipase A.** Spectra was obtained using Bruker instrument GN500 at 500 MHz, referenced to 4.79 ppm Deuterium Oxide peak. Integration analysis using TopSpin.

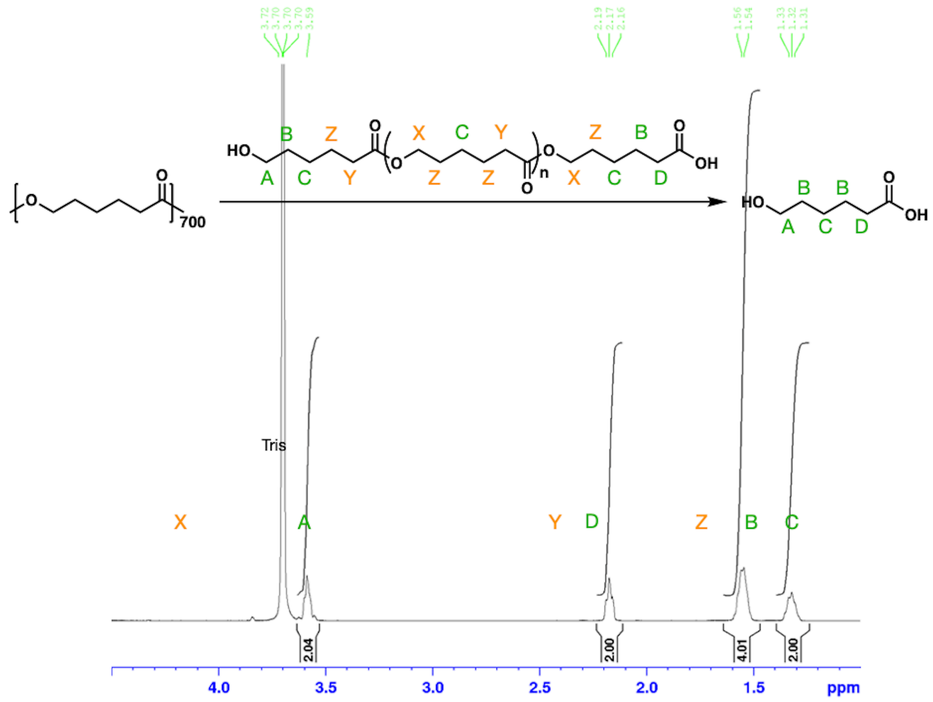


| C/D |
| --- |
| 1.00 |

**Supplementary Figure 7. [1]H NMR ($D_2O$) Spectra of the endpoint mixture collected from the PCL degradation process with free-floating enzyme Lipase A.** Spectra was obtained using Bruker instrument GN500 at 500 MHz, referenced to 4.79 ppm Deuterium Oxide peak. Integration analysis using TopSpin.

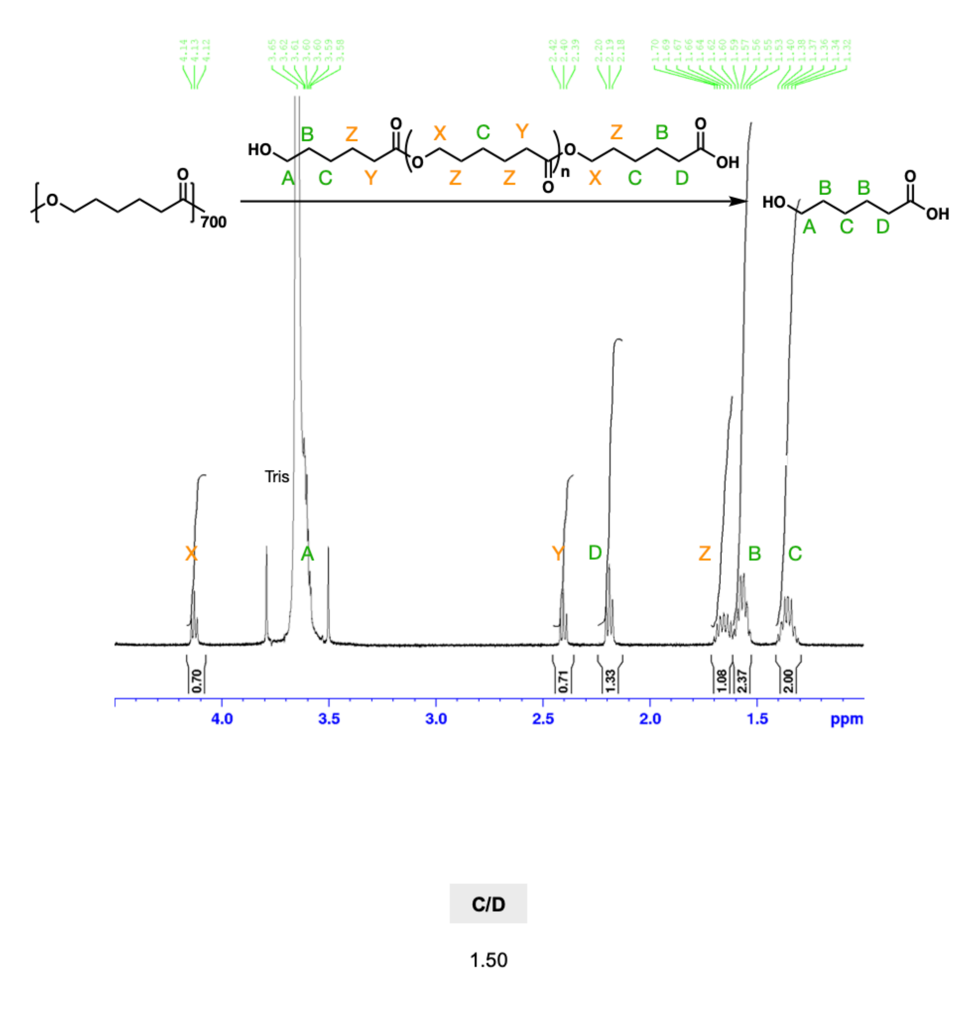


**Supplementary Figure 8. ¹H NMR ($D_2O$) Spectra of the midpoint mixture collected from the PCL degradation process with TIED-LipA.** Spectra was obtained using Bruker instrument GN500 at 500 MHz, referenced to 4.79 ppm Deuterium Oxide peak. Integration analysis using TopSpin.

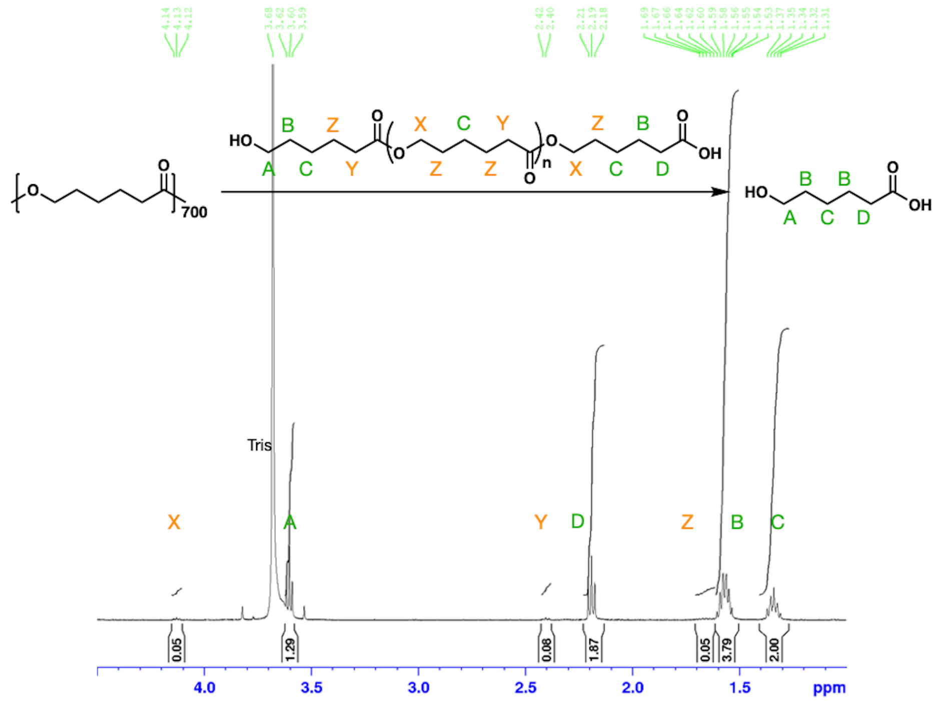


| C/D |
| --- |
| 1.07 |

**Supplementary Figure 9. $^1$H NMR ($D_2O$) Spectra of the endpoint mixture collected from the PCL degradation process with TIED-LipA.** Spectra was obtained using Bruker instrument GN500 at 500 MHz, referenced to 4.79 ppm Deuterium Oxide peak. Integration analysis using TopSpin.

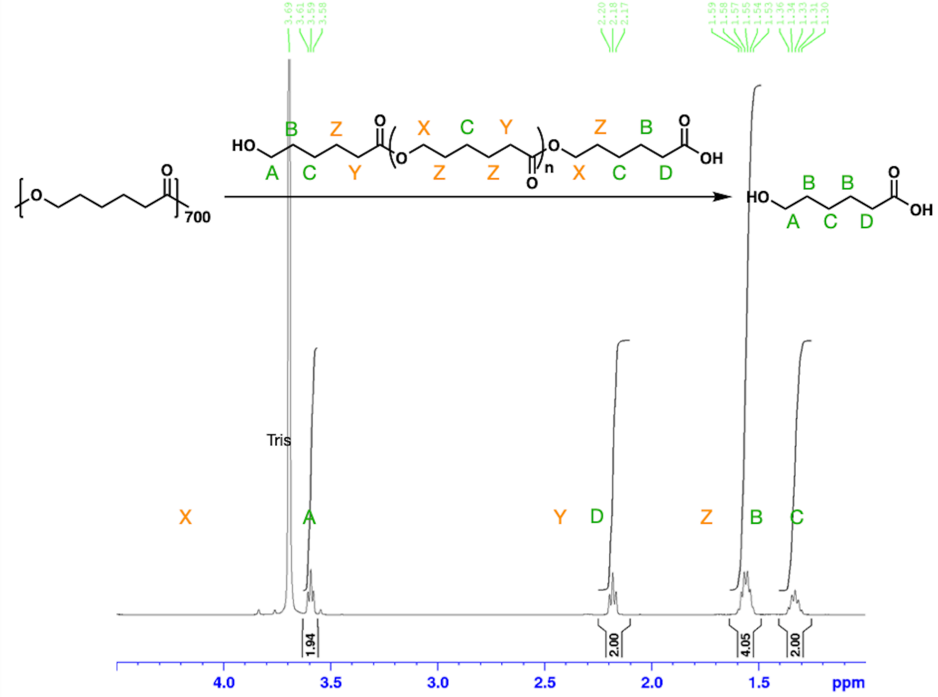


| C/D |
| --- |
| 1.00 |

**Supplementary Figure 10. [1]H NMR ($D_2O$) Spectra of the midpoint mixture collected from the PCL degradation process with free-floating enzyme Lipase B.** Spectra was obtained using Bruker instrument GN500 at 500 MHz, referenced to 4.79 ppm Deuterium Oxide peak. Integration analysis using TopSpin.

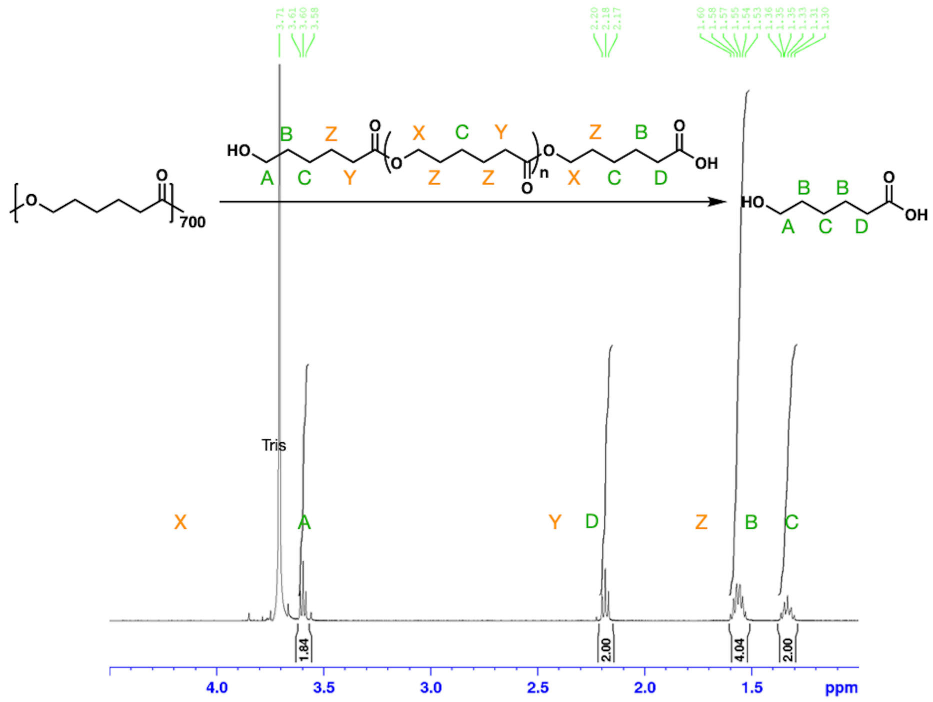


| C/D |
| --- |
| 1.00 |

**Supplementary Figure 11.** **[1]H NMR ($D_2O$) Spectra of the endpoint mixture collected from the PCL degradation process with free-floating enzyme Lipase B.** Spectra was obtained using Bruker instrument GN500 at 500 MHz, referenced to 4.79 ppm Deuterium Oxide peak. Integration analysis using TopSpin.

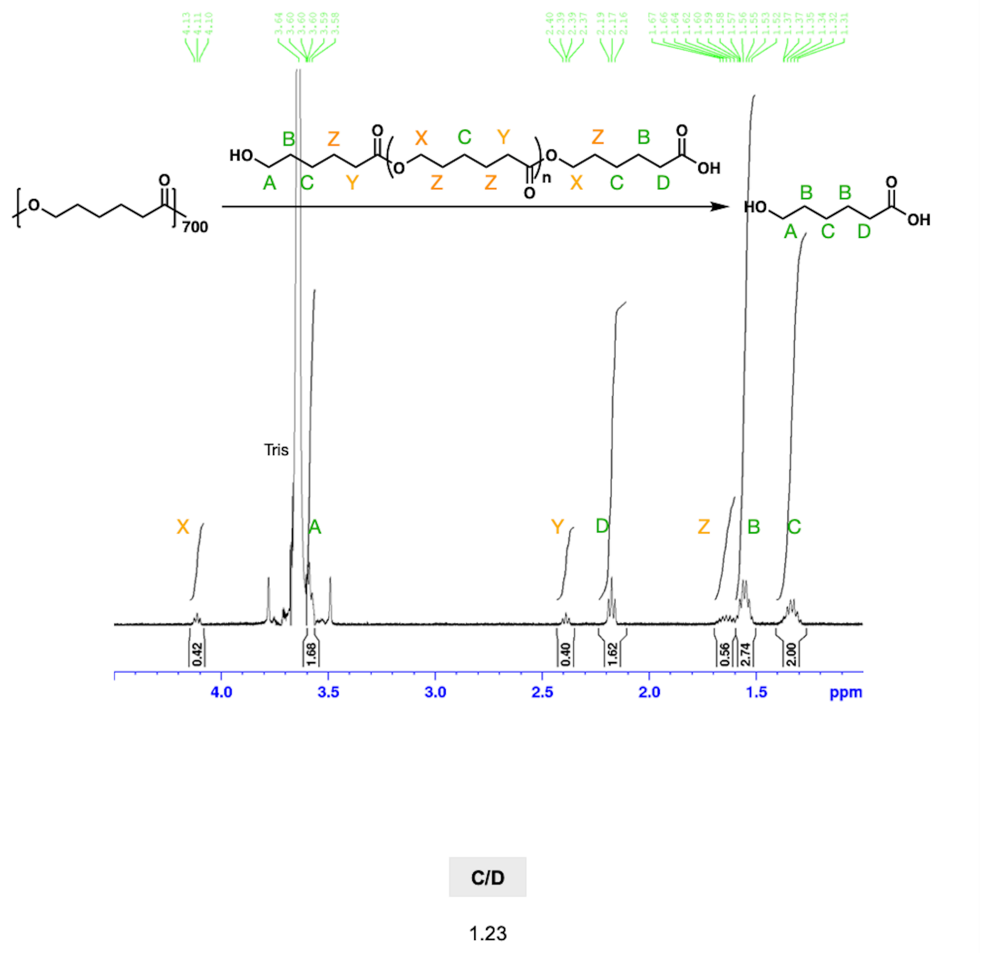


**Supplementary Figure 12. [1]H NMR ($D_2O$) Spectra of the midpoint mixture collected from the PCL degradation process with TIED-LipB.** Spectra was obtained using Bruker instrument GN500 at 500 MHz, referenced to 4.79 ppm Deuterium Oxide peak. Integration analysis using TopSpin.

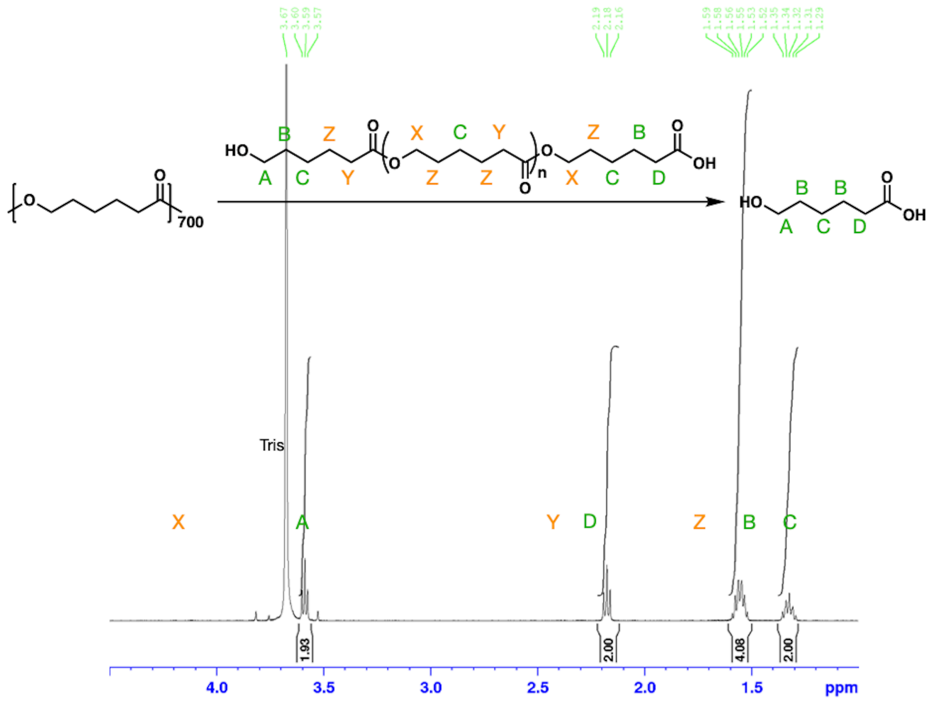


| C/D |
| --- |
| 1.00 |

**Supplementary Figure 13. [1]H NMR ($D_2O$) Spectra of the endpoint mixture collected from the PCL degradation process with TIED-LipB.** Spectra was obtained using Bruker instrument GN500 at 500 MHz, referenced to 4.79 ppm Deuterium Oxide peak. Integration analysis using TopSpin.

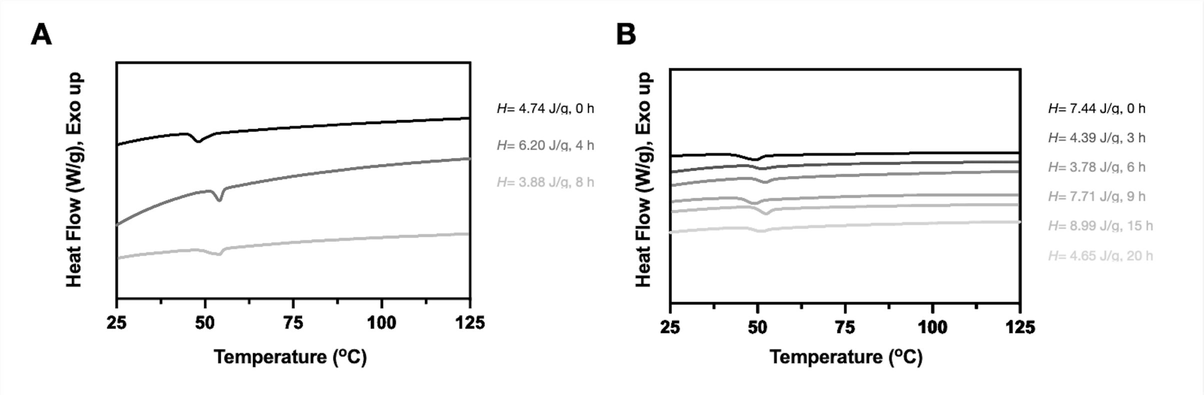


**Supplementary Figure 14. Differential scanning calorimetry (DSC) traces over time.** (A, B) Representative DSC traces of (A) PDLLA and (B) PLGA film degradation over time. Each trace is an individual film. The time mark shown in the figures represents how long they were incubated with TIED-LipA.

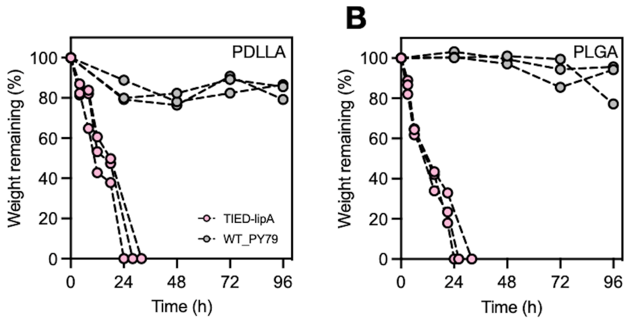


**Supplementary Figure 15. Differential scanning calorimetry (DSC) traces over time.** (A) Mass of the remaining PDLLA films upon incubation with 1 mL of TIED-LipA (pink) or wild-type spore (gray) suspension ($OD_{600}$ = 0.5). (B) Mass of the remaining PLGA films upon incubation with 1 mL of TIED-LipA (pink) or wild-type spore (gray) suspension ($OD_{600}$ = 0.5). Each circle represents individual mass data.

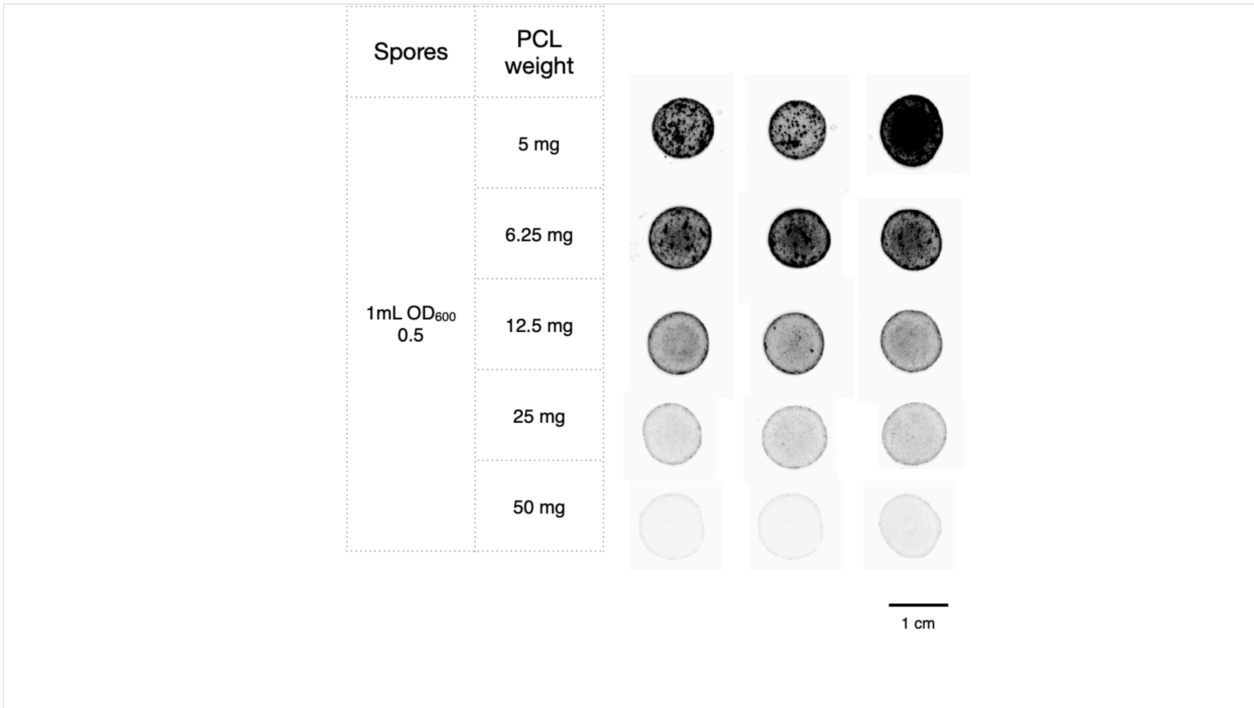


**Supplementary Figure 16. Biocomposite materials prepared with different spore loading.**

**Note:** Different weights of PCL pellets were dissolved in DCM at 3 w/v%. The solution was mixed with lyophilized fluorescent spores (TIED-mWasabi)[S1] in a powder form (from 1 mL of spore solution in $OD_{600}$= 0.5). 100 $\mu$L of the mixture was directly drop-cast into a film. The films were dried under ambient temperature and pressure overnight. The image was taken by Chemidoc MP (Bio-Rad), with Alexa 488 channel and 1 s exposure time. Based on this data, we concluded that a minimum amount of 12.5 mg PCL for 1 mL of spore solution in $OD_{600}$= 0.5 is necessary to ensure homogeneity. The biocomposite made with a higher PCL loading (25 mg) also showed complete degradation in the Tris-HCl buffer (Data not shown).

The PCL concentration in DCM (3 w/v%) was selected because lower concentrations than 2 w/v% had lower viscosity and could not hold the desired shape well in the drop-cast method. Higher concentrations than 4 w/v% was avoided due to spore dispersion issues and difficulties in processing the solution in the drop-cast method.

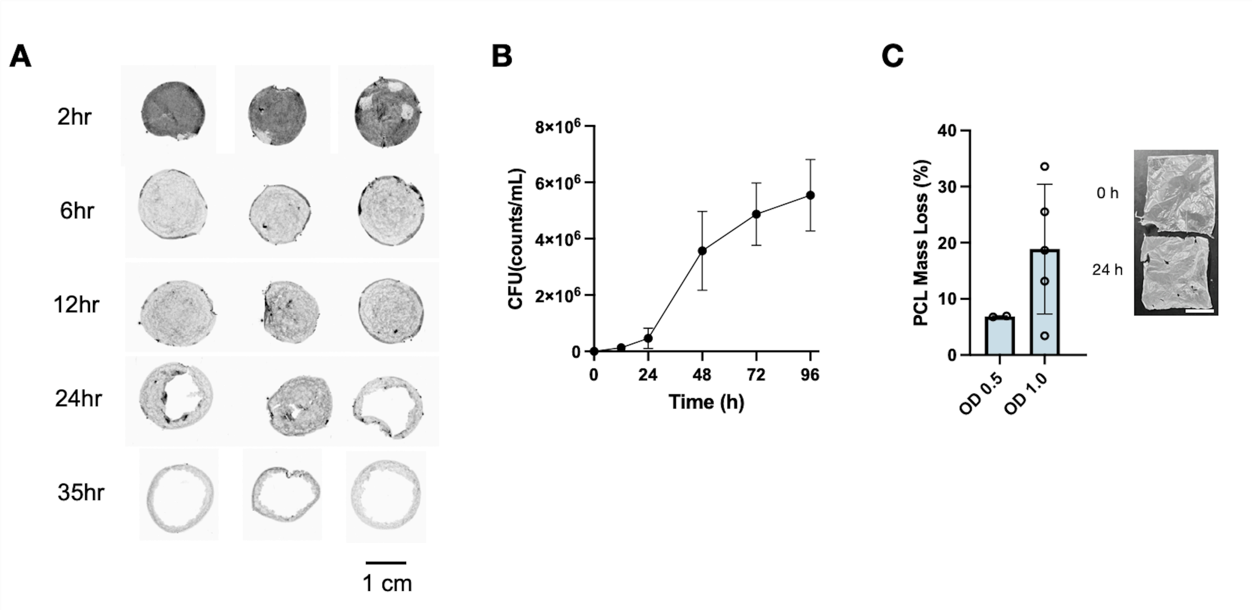


**Supplementary Figure 17. Spontaneous degradation of the biocomposite materials: materials' morphology and spores released from the materials.** (A) Images of degrading biocomposite materials. The images from top to bottom show the time progression. (B) Colony forming unit (CFU) from the supernatant collected from the degrading mixture. (C) The mass loss of additional PCL films treated with retrieved spores (left), and the morphology change in these films (right). Scale bar 1 cm.

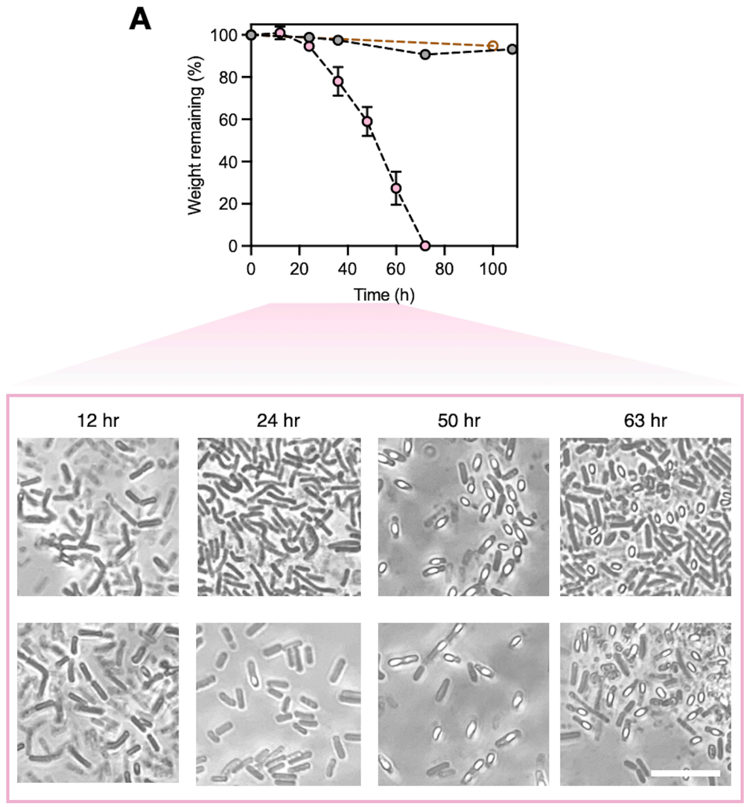


**Supplementary Figure 18. Degradation of biocomposite materials in LB medium.** (A) Mass loss of the biocomposite materials incorporating TIED-LipA (pink) and wild-type spore (gray) upon incubation in 1.5 mL LB medium. Mass loss of the PCL films incubated with TIED-LipA cells (brown) in 5 mL LB medium. (B) Optical microscopy images of the degradation mixture of biocomposite materials prepared with TIED-LipA (pink trace in A). Scale bar 10 $\mu$m.

### 3. Supplementary Tables

| Mn (kDa) | TIED-LipA | TIED-LipB | Free-floating LipA | Free-floating LipB |
|---|---|---|---|---|
| PCL (0 h) | **13.2** | | | |
| Midpoint (time) | **11.0** (2hr)<br>**13.7** (11hr)<br>**6.2** (17hr) | **13.0** (4hr)<br>**13.6** (15hr) | **13.3** (2hr) | **12.5** (0.4hr) |

**Supplementary Table 1. Number average molecular weight of all the PCL fragments calculated from GPC trace.**

| Mw (kDa) | TIED-LipA | TIED-LipB | Free-floating LipA | Free-floating LipB |
|---|---|---|---|---|
| PCL (0 h) | **20.5** | | | |
| Midpoint (time) | **17.2** (2hr)<br>**20.7** (11hr)<br>**9.5** (17hr) | **20.2** (4hr)<br>**20.8** (15hr) | **19.6** (2hr) | **19.8** (0.4hr) |

**Supplementary Table 2. Weight average molecular weight of all the PCL fragments calculated from GPC trace.**

| Dispersity | TIED-LipA | TIED-LipB | Free-floating LipA | Free-floating LipB |
|---|---|---|---|---|
| PCL (0 h) | **1.55** | | | |
| Midpoint (time) | **1.55** (2hr)<br>**1.51** (11hr)<br>**1.53** (17hr) | **1.55** (4hr)<br>**1.53** (15hr) | **1.47** (2hr) | **1.6** (0.4hr) |

**Supplementary Table 3. Dispersity of all the PCL fragments calculated from GPC trace.**

**Note:** Molecular weight numbers are acquired from a Polystyrene standard calibration curve.
